# Sensitive spatial genome wide expression profiling at cellular resolution

**DOI:** 10.1101/2020.03.12.989806

**Authors:** Robert R. Stickels, Evan Murray, Pawan Kumar, Jilong Li, Jamie L. Marshall, Daniela Di Bella, Paola Arlotta, Evan Z. Macosko, Fei Chen

## Abstract

The precise spatial localization of molecular signals within tissues richly informs the mechanisms of tissue formation and function. Previously, we developed Slide-seq, a technology which enables transcriptome-wide measurements with 10-micron spatial resolution. Here, we report new modifications to Slide-seq library generation, bead synthesis, and array indexing that markedly improve the mRNA capture sensitivity of the technology, approaching the efficiency of droplet-based single-cell RNAseq techniques. We demonstrate how this modified protocol, which we have termed Slide-seqV2, can be used effectively in biological contexts where high detection sensitivity is important. First, we deploy Slide-seqV2 to identify new dendritically localized mRNAs in the mouse hippocampus. Second, we integrate the spatial information of Slide-seq data with single-cell trajectory analysis tools to characterize the spatiotemporal development of the mouse neocortex. The combination of near-cellular resolution and high transcript detection will enable broad utility of Slide-seq across many experimental contexts.

## Main text

The *ab initio* identification of spatially defined gene expression patterns can provide powerful insights into the development and maintenance of complex tissue architectures, and the molecular characterization of pathological states. Recently, we developed Slide-seq^1^, a spatial genomics technology that quantifies expression genome-wide with high (10-micron) spatial resolution. Densely barcoded bead arrays, termed “pucks,” are fabricated by split-pool phosphoramidite synthesis, and indexed up front using a sequencing-by-ligation strategy. Downstream users of these arrays can perform assays with equipment found in a standard molecular biology laboratory, enabling the facile reconstruction of 3D tissue volumes that are tens or even hundreds of cubic millimeters in size.

However, Slide-seq’s low transcript detection sensitivity limited the range of biological problems to which the technology could be applied. Through improvements to the barcoded bead synthesis, the array sequencing pipeline, and the enzymatic processing of cDNA, we show here how we increased the sensitivity of Slide-seq by an order of magnitude. With our new protocol, termed Slide-seqV2, we demonstrate a range of new analytical possibilities by leveraging its improved capture efficiency, including the identification process-localized genes in neurons, and the analysis of developmental trajectories *in situ*.

We optimized the yield of Slide-seq capture by improving the array generation pipeline as well as the library preparation strategy (Fig. 1a). First, we developed a novel strategy to spatially index Slide-seq arrays using a monobase encoding scheme with sequencing by ligation using sequential interrogation by offset primer^2,3^ (Fig. S1, Methods). This strategy enables array indexing with open-source, commercially available reagents and increases the efficiency of spatial mapping of Illumina reads to bead barcodes by 50% (Fig. S1, mapped barcodes < hamming distance 2). In addition, we developed improved parameters for split-pool synthesis of the 10 μm polystyrene barcoded beads (Methods), which improved the clonality of our barcodes (Fig. S2). Together, these strategies enabled more efficient recovery of gene expression on Slide-seq arrays per Illumina read.

**Figure 1:**
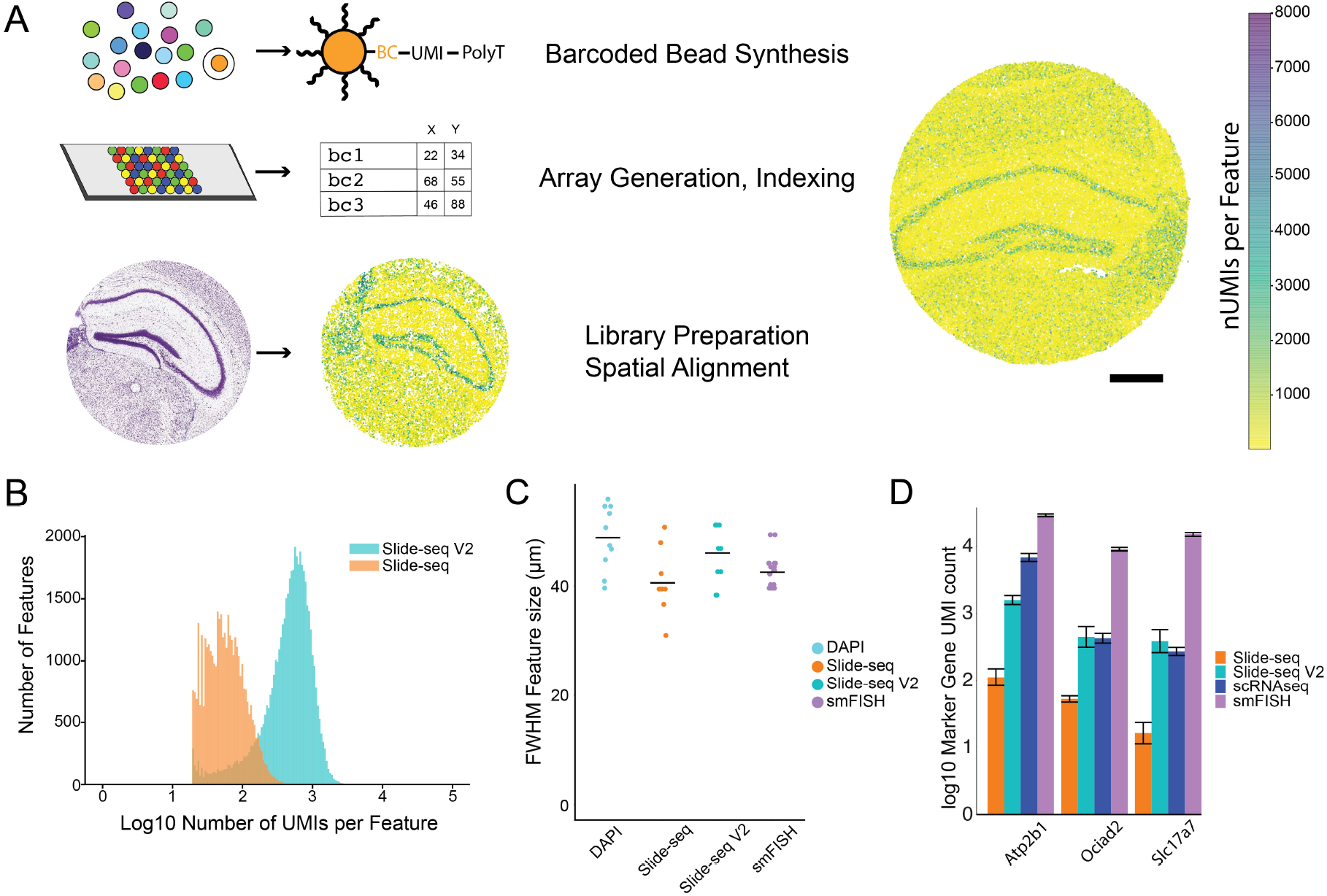
Highly improved mRNA detection sensitivity in Slide-seqV2. A) Overview of the Slide-seq method. Uniquely barcoded mRNA capture beads are affixed as a monolayer on a microscope slide, where their physical locations are determined by *in situ* sequencing. Subsequent application of a tissue section (mouse hippocampus, shown at bottom) to the array enables high-resolution spatial localization of gene expression. Right: An example array of mouse hippocampus generated with Slide-seqV2, with each bead colored by the number of UMIs. (scale bar 500 μm). B) Histogram of number of UMIs per bead for Slide-seq (red) versus Slide-seqV2 (blue) on serial mouse embryo sections. C) Measure of width of hippocampus CA1 across four modalities (N = 10 measurements per modality for DAPI (mean=48.8μm ∓ 5.8), Slide-seq(mean=40.6 μm ∓ 5.5), Slide-seqV2(45.9 ∓ 5.2). N=20 for smFISH (mean=42.5 μm ∓ 3.4). D) Comparison of marker gene counts in mouse hippocampus CA1 across four modalities (N = 6 measurements per modality, mean ∓ sd reported in Supp. Table 1, displayed in log10). For smFISH, Slide-seqV2 and Slide-seq data, all transcript counts within a fixed area of CA1 were summed together; for scRNA-seq, we summed the counts for the number of CA1 pyramidal cells counted within this area.

Next, we optimized the enzymatic library preparation steps of Slide-seq. We hypothesized that, due to the tissue’s inhibitory presence during reverse transcription, the template-switching reaction that adds a 3’ sequence priming site for whole-transcriptome amplification was inefficient. We therefore employed an additional second strand synthesis step^4^ after reverse transcription to increase the number of cDNAs that can be amplified by PCR. To evaluate the relative improvement in transcript capture efficiency, we performed Slide-seq on E12.5 mouse embryos. Using our improved protocol (Slide-seqV2), we obtained ~9.3x more transcripts (UMIs) per bead, compared to the original Slide-seq workflow (Fig. 1b, median 550 UMIs Slide-seqV2, 59 UMIs Slide-seq). In the mouse hippocampus, the capture efficiency of Slide-seqV2 was higher than that of a recently released commercial spatial transcriptomics technology^5^ (mean UMIs: Slide-seqV2 45,772, Visium = 27,952, for equal feature size, Fig. S3) while maintaining 25-100x improved spatial resolution (25x by area per feature, 100x including feature spacing, Fig. 1c).

Next, we sought to quantify the absolute sensitivity of Slide-seqV2 relative to other molecular technologies that measure RNA content in cells and tissues. We compared counts of known marker genes of mouse hippocampal CA1 pyramidal neurons (*Atp2b1, Ocaid2, Slc17a7)* in an equal number of pyramidal cells measured by: (1) Slide-seqV2; (2) Drop-seq, a high-throughput scRNA-seq method^6,7^; and (3) smFISH^8,9^ (Methods). We found that Slide-seqV2 detected only slightly fewer (17% fewer) counts than Drop-seq for the three genes measured (mean +/− std. scRNAseq = 33.5±1.4, 2.1±1.5,1.2±1.5, Slide-seqV2 = 15.7±1.5, 2.3±2.4, 1.9±2.6, Fig. 1d, N = 6, Supp. Table 1), demonstrating that Slide-seqV2 capture efficiency was competitive with modern single cell technologies.

We used Slide-seqV2 to gain insight into biological problems where higher capture sensitivity is important. Neurons actively transport specific mRNAs to dendrites and postsynaptic densities, where they play critical roles in synaptic development and plasticity^10–12^. Previous studies have explored dendritic enrichment through physical microdissection or cell culture, but none has systematically identified the distribution of dendritically localized transcripts *in situ*. Dendritic mRNAs constitute only a tiny fraction of neuronal transcripts^13^, necessitating higher sensitivity methods for their detection. To identify dendritically localized mRNAs from our mouse hippocampal Slide-seqV2 dataset, we took advantage of the stereotyped architecture of the CA1 neuropil to reduce the spatial localization of transcripts to a 1D profile perpendicular to the CA1 soma layer (from stratum oriens (s.o.) to stratum pyramidale (s.p.) across stratum radiatum (s.r.), Fig. 2a,b). For each gene detected in Slide-seq (N=4 sections), we calculated the spatial expression as a function of distance from the soma (representative spatial expression profiles shown in Fig. 2b, bottom).

**Figure 2:**
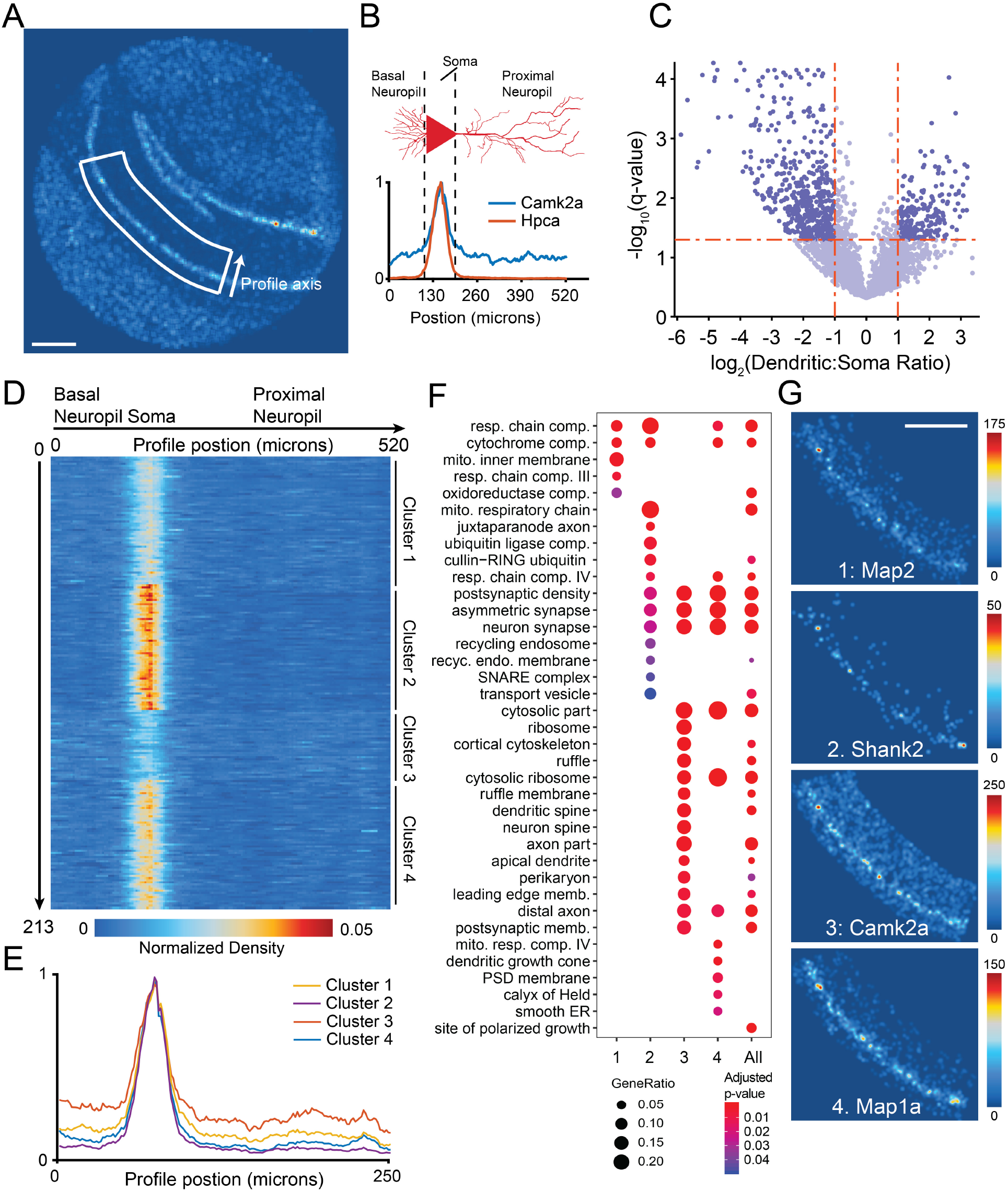
Slide-seq reveals spatial patterning of dendritically enriched mRNAs. A) Spatial heatmap of number of UMIs for a hippocampal Slide-seq dataset. B) (top) Schematic of linear spatial profiling across CA1 soma and dendrites. Line profiles perpendicular to the soma layer were averaged in the boxed region for each gene. (bottom) Spatial profiles of a CA1 marker (*Hpca*, red), and a classically dendritically localized gene (*Camk2a*, blue) are shown. C) Differentially expressed genes in soma versus proximal dendrites. Highlighted are genes with FDR-corrected p-value <0.05 and fold change >2 (Methods). D) Expression heatmap of 237 dendritically enriched RNAs. Columns move along profile position from 0 to 520 microns (each bin is 3 microns). Genes are shown clustered by their spatial profile (k-means clustering, 4 clusters). Rows are normalized and sum to 1. E) Average spatial expression profile of each of the four gene clusters identified in D across CA1. F) Gene-ontology classifications using over-representation analysis (Methods) for cellular component terms for each spatial cluster in **D** as well as all dendritically enriched genes. G) Slide-seq reconstruction images of one synaptic protein-encoding gene from each of the four clusters in **D**. Scale bars are 500 μm for all Slide-seq reconstructions.

To select for dendritically localized mRNA, we performed differential expression analysis, comparing the proximal neuropil (s.r.) to the soma (s.p.). The CA1 neuropil contains glial cell types (i.e. microglia and astrocytes) that also contribute RNA and interfere with analysis; we therefore included only genes expressed in CA1 pyramidal cells (>0.5 TPM in CA1 pyramidal neurons) and excluded those that are markers of non-neuronal cell types (Methods, Supp. Table 2), based on existing scRNA-seq data of the hippocampus^6^. After filtering, differential expression between the proximal neuropil and the soma revealed 213 significant genes with greater than 2-fold dendritic enrichment (Fig. 2c, unpaired t-test, N =4 sections, Supp. Table 2). These genes overlapped significantly (p<10^−16^, hypergeometric test, Fig. S4, Supp. Table 2) with two gene-lists of dendritically enriched RNAs from two previous studies^14,15^, suggesting Slide-seq can discover dendritically enriched genes.

Next, we asked whether functionally related genes showed similarities in their dendritic enrichment. First, we grouped dendritically enriched genes according to their 1D spatial expression profile (Fig. 2d). Using K-means, we identified 4 clusters of spatial expression of dendritically localized genes in CA1 neuropil, with clusters having different degrees of dendritic enrichment (Fig. 2e, Supp. Table 2). To identify whether this observed spatial diversity in localization was related to protein function, we used gene ontology (GO) to determine the cellular components of each spatial cluster (Fig. 2f, Methods). We found that each cluster was enriched for ontologically distinct groups of genes. Specifically, the first 2 clusters were enriched for components of the cellular respiration machinery, as well as ubiquitin ligases, while clusters 3 and 4 were enriched for ribosomal subunits. There was also a strong enrichment of synaptic proteins across 3 of the 4 clusters (2-4), suggesting that dendritically localized synaptic mRNAs demonstrate considerable heterogeneity in the degree of dendritic localization. Slide-seq’s genome-wide capture allowed us to visualize the heterogeneity in dendritic trafficking across two synaptic and two cytoskeletal genes chosen from each cluster (Fig. 2g, spatial reconstructions of all 213 genes are shown in Supp. Dataset 1). Taken together, these data demonstrate Slide-seqV2’s ability to characterize process-localized mRNAs, which appear to display significant heterogeneity amongst the various trafficked synaptic mRNA components.

During development, dynamic changes in gene expression across time and space help give rise to complex tissue architectures and terminally differentiated cell types. An array computational strategies have been developed to identify and explore developmental trajectories from scRNA-seq data^16–19^, based upon similarities in gene expression between individual profiles. More recently, an additional approach called RNA velocity was developed that dynamically models expression trajectories by the relative quantities of spliced and unspliced transcripts for each gene^20^. We reasoned that the combination of Slide-seqV2’s enhanced capture efficiency--which approaches that of scRNA-seq technologies--and its near-single-cell resolution--may allow us to exploit these powerful algorithms directly on our spatial data to learn how developmental processes are proceeding across a tissue section.

In the embryonic mouse neocortex, neuronal development progresses along a radial axis that begins in the Ventricular Zone (VZ) and moves through the Subventricular Zone (SVZ), Intermediate Zone (IZ), and finally the Cortical Plate (CP), where neurons integrate into cortical layers in a birthdate-dependent manner. We wondered whether Slide-seqV2 data could be used to successfully recover this highly spatially organized developmental trajectory^21^. We first applied unsupervised clustering^22^ to Slide-seqV2 data from embryonic day 15 (E15) developing mouse brain to characterize gene expression gradients in the neocortex. We annotated clusters corresponding to cell types in different developing brain regions, including cortex and striatum (Fig. 3a, Fig. S5). Segregating just the radially developing cortex (Fig. 3a, black box), we reclustered the beads to reveal populations representing the VZ, SVZ, IZ, CP, early cortical layers (L5/6) and Cajal Retzius cells (CR) (Fig. 3a).

**Figure 3:**
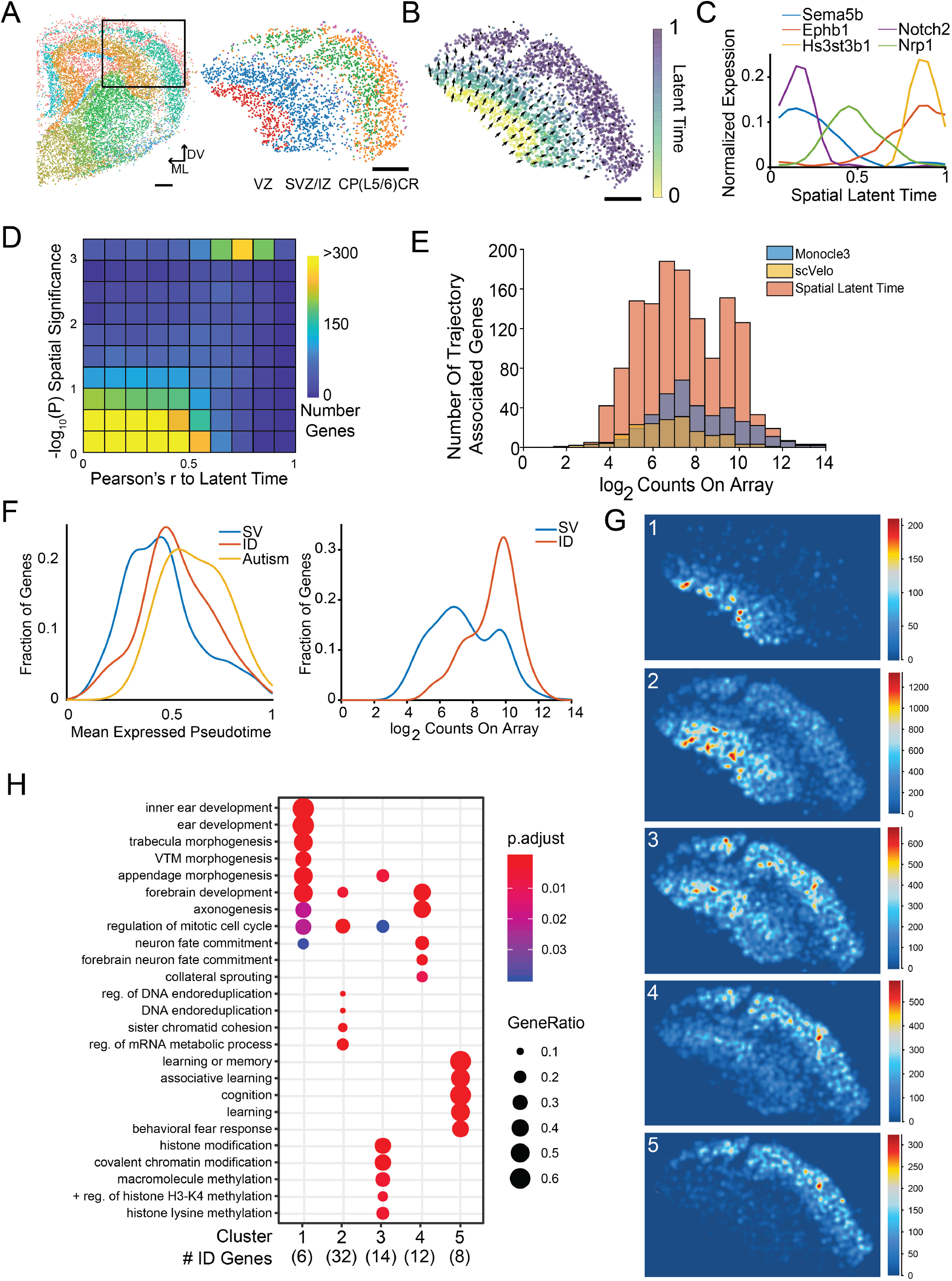
Slide-seq of developing mouse cortex reconstructs spatial developmental trajectories. A) Left: Unsupervised cluster analysis of Slide-seq data obtained from a section of E15 mouse brain. Black box indicates region chosen for downstream analysis. (Scale bar, 200 μm, ML: medial/lateral axis, DV: dorsal ventral axis). Right: Beads present within black-box inset from top, colored by their annotated cluster identities, subsetted by clusters of cortical identity. Red = Ventricular Zone (VZ), Blue/Purple = Subventricular Zone/ Intermediate Zone, Green/ Orange = Cortical Plate/ Layer 5 / 6, Pink = Cajal Retzius Cells (CR cells). B) Beads within the anatomical region of developing cortex, colored by their assigned latent time metric from scVelo. Arrow size and direction correspond to the direction and magnitude of the spatial derivative of the latent time in physical space. C) Expression profiles of sample genes jointly identified by Slide-seq, scVelo and Monocle3, across the Slide-seq-generated spatial latent time axis. D) Two-dimensional density plot quantifying the relationship between the a gene’s correlation with scVelo latent time (x-axis, Pearson’s r) and spatial significance (y-axis, log p-value, see Methods). Each square is colored by the number of genes found in that bin. E) Stacked histogram of the number of genes associated to the developmental trajectory by Monocle3 (blue), scVelo (yellow), and spatial latent time (red), binned by expression level (x-axis, log2 counts per gene across the dataset). F) left: density plot of all spatial latent time genes compared to DD latent time genes across mean expressed latent time value; right: density plot of all spatial latent time genes compared to DD latent time genes for summed gene expression across array. G) Slide-seq reconstruction images of metagenes associated with each spatial cluster of DD genes (Methods). H) Gene-ontology classifications using over-representation analysis (Methods) for biological process terms for each spatial cluster in **G.**

To determine whether Slide-seqV2 data can identify developmental trajectories, we first applied scVelo^23^, a recently developed trajectory inference method that leverages splicing information^20^, to order our beads along a predicted latent time (LT). Projection of each bead’s LT value onto spatial coordinates successfully recapitulated the established radial developmental axis of the neocortex (Fig. 3b). A very similar trajectory was recovered using the pseudotime ordering generated by Monocle3^17,24^(Fig. S5). Additionally, Slide-seqV2 data could also be used to recover the radial axis of ocular lens development in the embryonic eye (E12.5)^25^ (Fig. S5), demonstrating this analytical approach could be extended to other biological processes as well.

During the course of a developmental process, each stage of maturation can proceed at a different rate. We wondered whether Slide-seqV2’s spatial information could be exploited to identify the relative rates of differentiation across the radial axis of neocortical development. To accomplish this, we took the spatial derivative of the scVelo-generated LT (Methods), recovering regions where LT changes most dramatically (Fig. 3b with magnitude of arrows representing the magnitude of derivative). We found that the spatial rate of change was most pronounced at the earlier stages of the trajectory, decreasing as cells progress from VZ to SVZ/IZ, and largely terminating in the cortical plate.

Since each bead’s physical position is strongly predictive of its LT value, we reasoned that combining spatial and LT information could give us considerably greater statistical power to identify gene expression changes across this developmental process. The scVelo method was able to identify 179 genes with significant loading on LT, while the Monocle3 approach identified 377 genes. We previously demonstrated that we could leverage the spatial dimension of Slide-seq to systematically discover non-random spatial gene expression patterns^1^. Leveraging this, we identified 1349 spatially varying genes in the developing neocortex (P<0.005, Methods, Supp. Table 4), spatial expression plots of all genes are in Supp. Dataset 2). Among these were genes that are known to be involved in cortical development and are shared among Slide-seq and the trajectory inference methods including *Sema5b* and *Nrp1*, both involved in axonal guidance^26,27^ (Fig. 3c). We noted that these genes correlated strongly with the spatial LT axis. Thus, to systematically find genes that varied along this axis, we correlated the expression of these 1349 nonrandom genes with a spatial LT axis that was created by fitting a surface to the LT values in physical space (Methods). Of the 1349 spatially variable genes, 1043 correlated significantly with LT (pFdr <0.005), while very few of the non-spatially variable genes showed significant LT relationship (Fig. 3d). In addition, the 1043 genes were highly overlapping with the trajectory inference methods: amongst these were 76.5% of the scVelo-identified genes (137/179, Fig. S6, Supp. Table 3), and 75.6% of the Monocle3-defined genes (285/377, Fig. S6, Supp. Table 3). These results gave us confidence that the 1043 genes found using our spatial LT approach were truly associated with neocortical development.

Developmental disorders (DD) are a class of diseases frequently caused by pathogenic mutations in protein coding genes^28^ that often disrupt the normal process of neocortical development. Therefore, we asked how a set of 299 DD-associated genes that were recently discovered from exome sequencing of DD parent-offspring trios ^29^ distributed on our spatial LT trajectory. A total of 74 of the 299 DD-associated genes were found in the spatial LT gene set (1.87-fold enrichment, p = 3.2×10^−8^). These genes were expressed later in average LT compared with all spatial LT genes (Fig. 3f, left). Interestingly, the average expression of the 74 DD-associated spatial LT genes was much higher than all spatial LT genes (Fig. 3f, right). These 74 genes could be clustered into five groups based on their spatial expression patterns (Fig. 3g, Methods). The individual clusters were enriched for distinct GO functional terms, suggesting that these genes participate in distinct developmental processes and pathways (Fig. 3h), ranging from chromatin modification to establishment of neuronal states. Once additional phenotypic data becomes available about the relative clinical differences amongst these DD-associated genetic disorders, it will be revealing to understand how such phenotypes differentially load onto the spatial LT axis.

Here, we describe Slide-seqV2, an updated version of Slide-seq with nearly order of magnitude higher sensitivity. In particular, we demonstrated how the higher capture efficiency of Slide-seqV2 significantly expands the scope of possible analyses, including the discovery of genes with distinct patterns of subcellular localization, and the tracing of developmental trajectories through two-dimensional space. To further facilitate adoption of the technology, we have generated a streamlined pipeline for image processing and merging of short read sequencing and imaging data^30^ (Fig. S7, Methods). This pipeline provides statistics on the alignment of imaging and short read data, in addition to the gene expression matrix and spatial locations of each barcode, with limited user intervention. The combination of efficient molecular biology workflows, open sourced sequencing chemistry for array indexing, and easy-to-use software for merging imaging and sequencing data should support wide application of Slide-seq. We anticipate that the technical and computational improvements here will significantly accelerate the adoption of Slide-seq across the academic community.

## Acknowledgments

We thank Jordane Dimidschstein, and Gord Fishell for their helpful advice and suggestions on experimental analysis. This work was supported by an NIH New Innovator Award (DP2 AG058488-01 to E.Z.M.), an NIH Early Independence Award (DP5, 1DP5OD024583 to F.C.), the NHGRI (R01, R01HG010647 to E.Z.M. and F.C.), as well as the Schmidt Fellows Program at the Broad Institute and the Stanley Center for Psychiatric Research.

## Author Contributions

F.C. and E.Z.M. supervised the work. R.R.S. and E.M. performed experiments. R.R.S. and F.C analyzed the data. J.L. developed the Slide-seq tools software package. P.K. developed the bead synthesis protocol. J.L.M. performed optimization experiments. F.C., E.Z.M., R.R.S. and E.M. wrote the manuscript with input from all authors.

