## Supplemental_Information for "Sensitive spatial genome wide expression profiling at cellular resolution"

### Supplement

**Barcoded beads**

Bead barcodes were either synthesized by the Chemgenes Corporation or in-house on an Akta Oligopilot 10 on one of two polystyrene supports (Agilent PLRP-S-1000A 10 µm particles or 10 µm custom polystyrene from AMBiotech). Oligonucleotide synthesis was performed as described below. Beads were used with one of the two following sequences:

Chemgenes corporation beads:

5’-TTTTTTTTCTACACGACGCTCTTCCGATCTJJJJJJJTCTTCAGCGTTCCCGAGAJJJJJJJNNNNNNNNT30

Custom synthesis beads:

5'-TTT_PC_GCCGGTAATACGACTCACTATAGGGCTACACGACGCTCTTCCGATCTJJJJJJJTCTTCAGCGTTCCCGAGAJJJJJJTCNNNNNNNNT25 (vs1)

5'-TTT_PC_GCCGGTAATACGACTCACTATAGGGCTACACGACGCTCTTCCGATCTJJJJJJJTCTTCAGCGTTCCCGAGAJJJJJJNNNNNNNNVVT30 (vs2)

“PC” designates a photocleavable linker; “J” represents bases generated by split-pool barcoding, such that every oligo on a given bead has the same J bases; “N” represents bases generated by mixing, so every oligo on a given bead has different N bases; and “TX” represents a sequence of X thymidines. “V” represents bases which contain A,C,G and no T.

**Bead synthesis**

PLRP-S resin (~10 µm mean particle diameter) from Agilent Technologies were functionalized with a non-cleavable linker by Chemgenes Corp. The functionalized beads were then used as a solid support for reverse-direction phosphoramidite synthesis (5’ to 3’) on an Akta OligoPilot 10 using standard solid-phase DNA synthesis protocol. 5’-CE (b-cyanoethyl) phosphoramidites were purchased from Glen Research and were dissolved in anhydrous acetonitrile to obtain a concentration of 0.1M. Successive phosphoramidites were coupled for 5 minutes using 5-Benzylmercaptotetrazole (0.30 M in acetonitrile) as an activator. Oxidation of phosphite backbone to phosphate backbone was achieved using iodine. And failure sequences were capped using acetic anhydride. Dichloroacetic acid was used as detritylation reagent. For split-pool synthesis cycles, beads were suspended in acetonitrile and were divided into 4-equal portions. These bead aliquots were then placed in 4 separate synthesis columns and were reacted with either dG, dC, dT, or dA phosphoramidites. After each cycle, beads were pooled, suspended in acetonitrile and aliquoted into 4 equal portions. The split pool procedure was repeated 15 times in total (two blocks of 8 and 7 cycles) to obtain $4^{15}$ = 1,073,741,824 unique barcode sequences. After completion of the synthesis, the protecting groups from the nucleobases and phosphate backbone were removed by treating beads with 30% $NH_4OH$ containing 10% diethylamine for 40 h at room temperature. The beads were centrifuged and supernatant was discarded. Following this, beads were washed with 1% acetone in acetonitrile (3 times), water (3 times) and 10 mM tris buffer (pH = 8, 3 times).

**Puck Preparation**

Puck preparation was performed as described previously[1], with the following modification:

Beads were pelleted and resuspended water+ 10% DMSO at a concentration between 20,000 and 50,000 beads/µL , and 10µL of the resulting solution was pipetted into each position on the gasket. The coverslip-gasket filled with beads centrifuged at 40C, 850g for at least 30 minutes until the surface was dry.

**Puck Sequencing**

Puck sequencing was performed in a Bioptechs FCS2 flow cell using a RP-1 peristaltic pump (Rainin), and a modular valve positioner (Hamilton MVP). Flow rates between 1 mL/min and 3 mL/min were used during sequencing. Imaging was performed using a Nikon Eclipse Ti microscope with a Yokogawa CSU-W1 confocal scanner unit and an Andor Zyla 4.2 Plus camera. Images were acquired using a Nikon Plan Apo 10x/0.45 objective. After each ligation, images were acquired in the following channels: 488nm excitation with a 525/36 emission filter (MVI, 77074803); 561 nm excitation with a 582/15 emission filter (MVI, FF01-582/15-25); 561 nm excitation with a 624/40 emission filter (MVI, FF01-624/40-25); and 647 nm excitation with a 705/720 emission filter (MVI, 77074329). The final stitched images varied in size depending on the size of the Slide-seq array.

Pucks were sequenced using a sequencing-by-ligation approach, both with the SOLiD dibase-encoding strategy previously described[1,31] and with a monobase-encoding strategy developed for this work. Fluorescent oligonucleotides were synthesized on an Akta OligoPilot 10 or obtained from IDT (Supp. Table 5). A total of 8 fluorescent oligonucleotides were used and are referred to as 5(base) or 3(base) to indicate the corresponding mode of ligation and identity of the interrogated base.

The monobase strategy consisted of three ligation modes: 5' ligation, 3' ligation, and 5' ligation using SEDAL [2,32]. For both 5' and 3' ligation modes, a primer was injected into the flow cell at 5 µM concentration in 4x SASC for 40 minutes. Subsequently, the flow cell was washed in 5 mL of wash buffer (50 mM Tris-Acetate + 0.05% Triton-X 100). Ligation mix (recipes below) was then flowed into the chamber and allowed to sit for 40 minutes, at which point flow was reversed to return the ligation mix to its original reservoir. Ligation mix was reused for a complete sequencing run before being replenished. After a subsequent wash, pucks were imaged as described above and then stripped using 10 mL of 80% formamide for 20 minutes. For SEDAL ligations, a primer was added at 5 µM concentration to the ligation mix and this mixture was flowed into the chamber and allowed to sit for 2 hours.

Bead barcodes consisted of 15 "J" bases, of which 14 were used. In order to sequence these barcodes, we performed 3 rounds of SEDAL, 8 rounds of 5' ligation, and 3 rounds of 3' ligation (Fig. S1). The 14 primers necessary for this process were obtained from IDT (Supp. Table 5). The ligation mix recipes are given below:

5' Ligation mix:
1x T4 DNA Ligase Buffer (NEB)
6 U/µL T4 DNA Ligase (NEB)
20 µM each of 5T, 5A, 5G, and 5C oligonucleotides

3' Ligation mix:
1x T4 DNA Ligase Buffer (NEB)
6 U/µL T4 DNA Ligase (NEB)
20 µM each of 3T, 3A, 3G, and 3C oligonucleotides

SEDAL Ligation mix:
1x T4 DNA Ligase Buffer (NEB)
6 U/µL T4 DNA Ligase (NEB)
5 µM primer
5 µM each of 5T, 5A, 5G, and 5C oligonucleotides

**Microscopy**

Imaging was performed using a Nikon Eclipse Ti microscope with a Yokogawa CSU-W1 confocal scanner unit and an Andor Zyla 4.2 Plus camera. Images were acquired using a Nikon Plan Apo 10x/0.45 objective. After each ligation, we acquired four images: one using a 488 nm laser and a 525/36 emission filter (MVI, 77074803); one using a 561 nm laser and a 582/15 emission filter (MVI, FF01-582/15-25); one using a 561nm laser and a 624/40 emission filter (MVI, FF01-624/40-25); and one using a 647nm laser and a 705/72 emission filter (MVI, 77074329). The final stitched images were 6030 pixels by 6030 pixels.

**Image Processing and Basecalling**

Image processing was performed as previously described, and we have made an easy to use image processing and base calling matlab package that has been deposited into https://github.com/MacoskoLab/PuckCaller/. Input images are 4 channel sequencing images for each puck for each timepoint of sequencing. The outputs are, for each bead, a sequence string for the bead barcode. For monobase imaging the images are directly convertible to basespace rather than colorspace thus we omit the step of conversion of illumina reads to colorspace prior to comparison to the *in situ* indexing data as previously described.

##### ***Slide-seq tools***

We developed the Slide-seq tools for processing Slide-seq data. The workflow is illustrated in Fig. S7. The Slide-seq tools extract the barcode for each read in an Illumina lane from Illumina BCL files, collect, demultiplex, and sort reads across all of the tiles of a lane via barcode, and produce an unmapped bam file for each lane. The bam file is tagged with cellular barcode and molecular barcode. Low-quality reads are filtered, and reads are trimmed with starting sequence and adapter-aware poly A. STAR is used to align reads to genome sequence. The unmapped bam and sorted aligned bam are merged and tagged with interval and gene function. Top cells are selected that a number of transcripts are aligned to. For each of those barcodes with short read sequencing, hamming distances are calculated between it and all of bead barcodes from in situ sequencing. The list of unique matched Illumina barcodes with hamming distance <= 1 along with the matched bead barcodes are outputted. Finally a few reports and plots are generated on the alignment outputs and the matched bead barcodes, such as read quality and mapping rate plot, digital expression matrix, color-scaled number of UMIs per bead, etc. Drop-seq tools, Picard and Samtools are called in the Slide-seq tools. The scripts, documentations, and example data are available at https://github.com/MacoskoLab/slideseq-tools.

Supplementary Table 7 shows running time of the Slide-seq tools on four libraries: 190926_01, 190926_02, 190926_03 and 190926_06. The Illumina platform is NovaSeq, and there are two lanes in the experiment. Reads were aligned to GRCm38.81 genome sequence. The read base quality for alignment was set to 10. The minimum number of transcripts per cell for selecting top cells was set to 10 as well. Reads aligned to both exons and introns were involved in the gene expression analysis. In order to speed up the process, the Slide-seq tools split each lane of NovaSeq data into 10 slices, parallel ran the alignment steps on the slices and combined the alignment outputs together.

##### **Slide-seqV2 library preparation**

RNA Hybridization:

Pucks in 1.5 mL tubes were immersed in 200 μL of hybridization buffer (6x SSC with 2 U/μL Lucigen NxGen RNAse inhibitor) for 30 minutes at room temperature to allow for binding of the RNA to the oligos on the beads.

First Strand Synthesis
Subsequently, first strand synthesis was performed by incubating the pucks in RT solution for 1.5 hours at 52 C.

RT solution:

- 115 µL H2O
- 40 µL Maxima 5x RT Buffer (Thermofisher, EP0751)
- 20 µL 10 mM dNTPs (NEB N0477L)
- 5 µL RNase Inhibitor (Lucigen 30281)
- 10 µL 50 µM Template Switch Oligo (Qiagen #339414YCO0076714)
- 10 µL Maxima H- RTase (Thermofisher, EP0751)

#### Tissue Digestion:

200 µL of 2x tissue digestion buffer was then added directly to the RT solution and the mixture was incubated at 37C for 30 minutes.

2x tissue digestion buffer:

200 mM Tris-Cl pH 8
400 mM NaCl
4% SDS
10 mM EDTA
32 U/mL Proteinase K (NEB P8107S)

Second Strand Synthesis:

The solution was then pipetted up and down vigorously to remove beads from the surface, and the glass substrate was removed from the tube using forceps and discarded. 200 µL of Wash Buffer was then added to the 400 µL of tissue clearing and RT solution mix and the tube was then centrifuged for 3 minutes at 3000 RCF. The supernatant was then removed from the bead pellet, the beads were resuspended in 200 µL of Wash Buffer, and were centrifuged again. This was repeated a total of three times. The supernatant was then removed from the pellet. The beads were then resuspended in 200µl of ExoI mix and incubated at 37 degrees for 50mins.

Wash Buffer:

- 10 mM Tris pH 8.0
- 1 mM EDTA
- 0.01% Tween-20

ExoI mix:

- 170µl H20
- 20µl ExoI buffer
- 10µl ExoI (NEB M0568)

After ExoI treatment the beads were centrifuged for 3 minutes at 3000 RCF. The supernatant was then removed from the bead pellet, the beads were resuspended in 200 µL of Wash Buffer, and were centrifuged again. This was repeated a total of three times. The supernatant was then removed from the pellet. The pellet was then resuspended in 200µl of 0.1 N NaOH and incubated for 5 minutes at room temp. To quench the reaction 200µl of Wash Buffer was added and beads were centrifuged for 3 minutes at 3000 RCF. The supernatant was then removed from the bead pellet, the beads were resuspended in 200 µL of Wash Buffer, and were centrifuged again. This was repeated a total of three times. Second Strand Synthesis was then performed on the beads by incubating the pellet in 200µ of Second Strand Mix at 37 degrees for 1 hour.

Second Strand Synthesis mix:

- 133 µl H2O
- 40 µl Maxima 5x RT Buffer
- 20 µl 10 mM dNTPs
- 2µl 1mM dN-SMRT oligo
- 5µl Klenow Enzyme (NEB M0210)

After Second Strand Synthesis 200µl of Wash Buffer was added and the beads were centrifuged for 3 minutes at 3000 RCF. The supernatant was then removed from the bead pellet, the beads were resuspended in 200 µL of Wash Buffer, and were centrifuged again. This was repeated a total of three times.

Library Amplification:

200µl of water was then added to the bead pellet and the beads were moved into a 200 µL PCR strip tube, pelleted in a minifuge, and resuspended in 200 µL of water. The beads were then pelleted and resuspended in library PCR mix and PCR was performed as outlined below:

Library PCR mix:

- 22 µL H2O
- 25 µL of Terra Direct PCR mix Buffer (Takara Biosciences 639270)
- 1µl of Terra Polymerase (Takara Biosciences 639270)
- 1 µL of 100 µM Truseq PCR handle primer (IDT)
- 1 µL of 100 µM SMART PCR primer (IDT)

PCR program:

- 95 C 3 minutes
- 4 cycles of:
  - 98 C 20 s
  - 65 C 45 s
  - 72 C 3 min
- 9 cycles of:
  - 98 C 20 s
  - 67 C 20
  - 72 C 3 min
- Then:
  - 72 C 5 min
  - 4 C forever

#### PCR cleanup and Nextera Tagmentation:

The PCR product was then purified by adding 30 µL of Ampure XP (Beckman Coulter A63880) beads to 50 µL of PCR product. The samples were cleaned according to manufacturer's instructions and resuspended into 50 µL of water and the cleanup was repeated resuspending in a final concentration of 10µl. 1 µL of the library was quantified on an Agilent Bioanalyzer High sensitivity DNA chip (Agilent 5067-4626). Then, 600 pg of PCR product was taken from the PCR product and prepared into Illumina sequencing libraries through tagmentation with Nextera XT kit (Illumina FC-131-1096). Tagmentation was performed according to manufacturer's instructions and the library was amplified with primers Truseq5 and N700 series barcoded index primers. The PCR program was as follows:

72°C for 3 minutes
95°C for 30 seconds
12 cycles of:
    95°C for 10 seconds
    55°C for 30 seconds
    72°C for 30 seconds
72°C for 5 minutes
Hold at 10°C

Samples were cleaned with AMPURE XP (Beckman Coulter A63880) beads in accordance with manufacturer's instructions at a 0.6x bead/sample ratio (30 µL of beads to 50 µL of sample) and resuspended in 10µL of water. Library quantification was performed using the Bioanalyzer. Finally, the library concentration was normalized to 4nM for sequencing. Samples were sequenced on the Illumina NovaSeq S2 flowcell 100 cycle kit with 12 samples per run (6 samples per lane) with the read structure 42 bases Read 1, 8 bases i7 index read, 50 bases Read 2. Each puck received approximately 200-400 million reads, corresponding to 3,000-5,000 reads per bead.

**Animal Handling**

All procedures involving animals at the Broad Institute were conducted in accordance with the US National Institutes of Health Guide for the Care and Use of Laboratory Animals under protocol number 0120-09-16.

**Transcardial Perfusion**

Animals were anesthetized by administration of isoflurane in a gas chamber flowing 3% isoflurane for 1 minute. Anesthesia was confirmed by checking for a negative tail pinch response. Animals were moved to a dissection tray and anesthesia was prolonged via a nose cone flowing 3% isoflurane for the duration of the procedure. Transcardial perfusions were performed with ice cold pH 7.4 HEPES buffer containing 110 mM NaCl, 10 mM HEPES, 25 mM glucose, 75 mM sucrose, 7.5 mM $MgCl_2$, and 2.5 mM KCl to remove blood from brain and other organs sampled. The appropriate organs were removed and frozen for 3 minutes in liquid nitrogen vapor and moved to -80C for long term storage.

**Tissue Handling**

Fresh frozen tissue was warmed to -20 C in a cryostat (Leica CM3050S) for 20 minutes prior to handling. Tissue was then mounted onto a cutting block with OCT and sliced at a 5° cutting angle at 10 µm thickness. Pucks were then placed on the cutting stage and tissue was maneuvered onto the pucks. The tissue was then melted onto the puck by moving the puck off the stage and placing a finger on the bottom side of the glass. The puck was then removed from the cryostat and placed into a 1.5 mL eppendorf tube. The sample library was then prepared as below. The remaining tissue was re-deposited at -80 C and stored for processing at a later date.

**Diffusion Analysis**

Determination of diffusion was determined as previously described by measuring features across CA1 mouse hippocampus[1]. This time Slide-seqV2 data was included in addition to data obtained from smFISH, DAPI staining, and Slide-seq.

**Comparison of counts for Slide-seq, Slide-seqV2, FISH, and scRNAseq**

For Slide-seq we subsetted a region of CA1 and took the total number of counts for each of the marker genes. A serial section was stained with DAPI and segmented in ImageJ by first scaling signal to background and binarizing the image followed by applying a 1.5 µm Gaussian Blur and a watershed transform. A box was taken matching the size of that taken in the Slide-seq data and the total number of nuclei were counted that had a diameter greater than 2µm and less than 12 µm. The total counts for each of the genes obtained from Slide-seq data was then divided by the number of nuclei present in that same area. The same methodology was performed for Slide-seqV2.

For the scRNAseq data, we used the mouse brain scRNAseq data from Saunders et al 2018 and pulled an equal number of cells found from the Slide-seq data from the cluster representing the hippocampal CA1 neurons.

For the smFISH data we generated the data by using HCRV3.0 with probes sets against each of the genes chosen in 488nm, 555nm, and 594nm. We stained the tissue with DAPI for segmentation purposes and for counting nuclei. We performed nuclei identification and counting as described above and performed counts in the FISH data over the same area using StarSearch.

**Comparison of Slide-seq to 10x Visium technology**

To compare Slide-seq to 10x Visium data [5] we downloaded available coronal mouse hippocampus data and plotted the number of UMIs per spatial feature. We next binned

Slide-seq data from the same region to equivalent feature size (110uμm), merging the counts of Slide-seq beads (10μm original) within each of the larger features generated (110μm). We then plotted the binned Slide-seq data alongside the same region in Visium with equal scaling for number of UMIs. Scale bars in each image represent number of UMIs. We also show a histogram of number of UMIs recovered per feature for the Slide-seqV2 with 100μm feature size and for the 10x visium data with the dotted line representative of the mean for each dataset.

**Hippocampal Slide-seq**

Slide-seqV2 was performed on the mouse hippocampus (N=4 sections, 2 mice). A spline was fit along the pyramidal cells layer of CA1. Beads were averaged to a profile perpendicular to this spline ~100 micrometers into the Basal neuropil, and ~400 microns to the proximal neuropil to form a spatial profile of gene expression along the CA-1 neuropil axis.

**Dendritic enrichment analysis**

To test for dendritic enrichment, for each gene, the gene expression in the soma layer (defined as +/- 32.5 micron from the peak of the profile counts for all genes) was compared against the gene expression in the proximal dendrites (greater than 32.5 micron away from the peak of the CA1 layer). For each gene, the gene expression in the Soma layer was normalized to the total number of UMI-counts in the Soma layer, and the gene expression in the proximal dendrite layer was normalized to the total number of UMI-counts in the proximal dendrite layer. A two sample t-test was performed to identify differentially expressed genes, and pFDR was calculated as described previously[33].

**Spatial clustering of dendritically enriched genes**

For the 213 genes identified to be dendritically enriched, we clustered genes by their spatial profile along the CA1-neuropil axis via k-means clustering. The gap-statistic was used to determine the optimal number of clusters (4).

**GO analysis**

For each cluster identified by spatial profiling, GO analysis was performed using the clusterProfiler [34]package in R. Cellular components from the org.Mm.eg.db[35]:. Genome wide annotation for Mouse in Bioconductor was used for the ontology database. For Fig. 3, GO analysis was performed using Biological processes from org.Mm.eg.db.

**Embryo samples**

Whole mount frozen embryos were obtained from a commercial source Zyagen (San Diego, CA). The pregnant mice (C57BL/6NCrl) were bred and maintained by Charles River Laboratories. The time-pregnant mice (day 10 or day 12) were shipped to Zyagen (San Diego, CA) the same day. The mice were sacrificed on the day of arrival for embryo collection.

**Trajectory analysis**

Trajectory analysis was performed using the recently released method scVelo[23]. We first loaded intronic and exonic gene expression matrices, UMAP coordinates created in Seurat from the original clustering of the Slide-seq data, cluster IDs, and spatial coordinates of each bead from Slide-seq into a scanpy object using a custom python environment. We next applied the latent time method developed in scVelo to our Slide-seq expression data and plotted each bead using the Slide-seq coordinates with the shading defined by the latent time ordering. Plots of individual expression of genes over latent time were generated using plotting functions in scVelo plotting the expression of each individual gene over the latent time axis with coloring of each bead by cluster identity to the original clustering of the data. Plots showing expression for each of the genes on the puck were performed using a custom python script. The gene lists for latent time were called as velocity loading genes from the scVelo pipeline using standard parameters and a likelihood cutoff of >0.1.

Monocle3 [17,24] was run on the data by importing the UMAP and PCA coordinates from Seurat into a Single Cell Experiment object. The analysis was performed in accordance with the monocle tutorial found at. http://cole-trapnell-lab.github.io/monocle-release/monocle3/ . The q-value cutoff for gene selection was q<0.005.

**Fitting a spatial surface to latent time**

Latent time data scores generated from scVelo and spatial coordinates were taken as a 3D set of points (x,y, latent time score) and a surface was fit over the set of points for a region of the cortex. A grid was created (80μm x 80m for cortex) and the spatial derivative was taken over the grid using matlab's differentiate function. The fx, fy of the surface were extracted from matlab and imported into python. The plot for Fig. 3b was generated using a custom python script where the magnitude of the arrows represents the magnitude of the derivative at each of the points in the grid. The position of the underlying beads is from Slide-seq and the color is from scVelo's latent time output.

**Spatially non-random gene analysis:**

Test for spatial non-randomness was performed as previously described [1] with the following modifications:

Genes were identified as spatially non-random using a custom Matlab application. The set of pairwise Euclidean distances between all beads was calculated. Candidate genes for the statistical significance analysis were required to have at least one transcript on at least 10 beads. To determine whether a transcript had a significantly non-random spatial distribution within a particular set of beads, we compared the distribution of pairwise distances between the beads expressing at least one count of that transcript to the distribution of pairwise distances between an identical number of beads, sampled randomly from all mapped beads on the puck with probability proportional to the total number of transcripts on the bead. Specifically, we generated 1000 such random samples, and for each sample calculated the distribution of pairwise distances. We then calculated the average distribution of pairwise distances, averages pairwise across all 1000 samples. Finally, we calculated the L1 norm between the distribution of pairwise distances for each of the 1000 random samples and the average distribution, and the L1 norm between the distribution of pairwise distances for the true sample of beads and the average distribution. We defined *p* to be the fraction of random samples having distributions closer to the average distribution (under the L1 norm) than the true sample, and considered any genes with values $p<=0.005$.

**Spatial correlation to latent time:**

Spatially identified genes were binned along 20 spatial contours of the same latent time as fitted by the surface described above. Expression was normalized for each bin by the total number of counts observed. For each gene, the Pearson's correlation coefficient and the pvalue of the correlation between the binned expression in the spatial latent time axis was correlated with a linear function of slope 1. pFDR was calculated as described previously[33].

**Spatial clustering of DD genes**

For the 74 genes identified to be involved in developmental disorders that load onto pseudotime, the spatial correlation between each gene was determined by convolving the spatial expression of each gene with an integralBoxFilter of size 70 microns, and then the 2-D cross-correlation for each gene against each other gene was calculated with the matlab function corr2. The spatial cross-correlation matrix was clustered using k-means, the gap-statistic was used to determine the optimal number of clusters (6).

**Supplemental Dataset 1:**Plots of all genes dendritically enriched in Slide-seq

**Supplemental Dataset 2:** Plots of all genes called as Spatially Significant in Slide-seq

**Supplementary Table 1**: Mean and Standard Deviation of Gene Measurements Fig. 1c (total over 200 cells).

| | smFISH | scRNAseq | Slide-seq | Slide-seqV2 |
|---|---|---|---|---|
| Atp2b1 | 28480∓295 | 6700∓225 | 133∓17 | 3131∓73 |
| Ociad2 | 9016∓139 | 424∓22 | 53.5∓6 | 444∓17 |
| Slc17a7 | 14928∓225 | 246∓9 | 21.5∓3 | 383∓12 |

**Supplementary Table 2**: Dendritically enriched gene-sets.

Sheet 1: Dendritically enriched RNAs identified by slide-seq along with Fold-Change enrichment, and FDR corrected q-value.

Sheet 2: Grouping of dendritically enriched genes by spatial clusters.

Sheet 3: List of genes which overlap with Tushev et al., 2018, and Ainsley et al., 2016 as well as genes uniquely identified by Slide-seq.

Sheet 4: Genes taken over latent time along with likelihood

**Supplementary Table 3**: List of genes unique to each method regarding the trajectory inference:

Sheet 1: Genes unique to Slide-seq
Sheet 2: Genes unique to monocle3
Sheet 3: Genes unique to scVelo

**Supplementary Table 4**: List of all genes called as Spatially Significant for Slide-seq data in the cortex

**Supplementary Table 5**: Oligonucleotide sequences used in the study.

| Name | Sequence |
|---|---|
| Template Switch Oligo | AAGCAGTGGTATCAACGCAGAGTGAATrG+GrG |
| Truseq_PCR_Handle | CTACACGACGCTCTTCCGATCT |
| SMART_PCR_Primer | AAGCAGTGGTATCAACGCAGAGT |
| dN-SMRT oligo | AAGCAGTGGTATCAACGCAGAGTGANNNGGNNNB |
| Truseq P5 | AATGATACGGCGACCACCGAGATCTACACTCTTTCCCTACACGACGCTCTTCCGATCT |
| Truseq-1 | /5phos/GATCGGAAGAGCGTCGTGTAG |
| Truseq | /5phos/AGATCGGAAGAGCGTCGTGTAG |
| Truseq+1 | /5phos/NAGATCGGAAGAGCGTCGTGTAG |
| Truseq+2 | /5phos/NNAGATCGGAAGAGCGTCGTGTAG |
| Truseq(8b)+3 | /5phos/NNNAGATCGGA/3InvdT/ |

| | |
|---|---|
| 3UP-1 | TCTCGGGAACGCTGAAG |
| 3UP | TCTCGGGAACGCTGAAGA |
| 3UP+1 | TCTCGGGAACGCTGAAGAN |
| UP-1 | /5phos/CTCGGGAACGCTGAAGA |
| UP | /5phos/TCTCGGGAACGCTGAAGA |
| UP+1 | /5phos/NTCTCGGGAACGCTGAAGA |
| UP+2 | /5phos/NNTCTCGGGAACGCTGAAGA |
| UP(7b)+3 | /5phos/NNNTCTCGGG/3InvdT/ |
| UP(7b)+4 | /5phos/NNNNTCTCGGG/3InvdT/ |
| Monobase 5A | /6FAM/IIINNNAN |
| Monobase 5G | /aquaphluor593/IIINNNGN |
| Monobase 5C | /CY3/IIINNNCN |
| Monobase 5T | /Cy5/IIINNNTN |
| Monobase 3A | /5Phos/NANNNIII/6FAM/ |
| Monobase 3G | /5Phos/NGNNNIII/aquaphluor593/ |
| Monobase 3C | /5Phos/NCNNNIII/Cy3/ |
| Monobase 3T | /5Phos/NTNNNIII/Cy5/ |

**Supplementary Table 6**: Slide-seq pucks used in each experiment.

| Figure | Puck Used (Tissue Type) |
|---|---|
| 1A | Left, Puck_190921_21 (mouse hippocampus), right, Puck_200115_08 (mouse hippocampus) |

| | |
|---|---|
| 1B | Puck_190926_03 (mouse embryo Slide-seqV2), Puck 191007_07 (mouse embryo Slide-seq) |
| 1C/D | Puck_190921_21(mouse hippocampus, Slide-seqV2) |
| 2A-G | Puck_191204_01 (mouse hippocampus) |
| 3 | Puck_190921_19 (mouse E15 brain) |
| S3 | Puck_200115_08 (mouse hippocampus), visium data is coronal section from 10x website |
| S4 | Puck_191204_01 (mouse hippocampus) |
| S6 | Puck_190921_19 (mouse E15 brain) |
| Supplementary Table 7 | Puck_190926_01, Puck_190926_02, Puck_190926_03, Puck_190926_06 |

**Supplementary Table 7**: Running time of the Slide-seq tools pipeline.

| Steps | # libraries | # lanes | # slices | Parallel | Size of bam | # barcodes | Time (minutes) |
|---|---|---|---|---|---|---|---|
| Extract Illumina barcodes | 4 | 1 | 10 | Per lane | | | 44 |
| Convert Illumina basecalls to bam | 4 | 1 | 1 | Per slice | 600M | | 25 |
| Align reads | 1 | 1 | 1 | Per slice | 600M | | 35 |
| Merge and validate bam | 1 | 2 | 10 | | 10G | | 60 |

| | | | | | | | |
|---|---|---|---|---|---|---|---|
| Select cells by the number of transcripts | 1 | 2 | 10 | | 10G | 300K | 50 |
| Generate alignment reports and plots | 1 | 2 | 10 | | 10G | | 70 |
| Barcode matching | 1 | 2 | 10 | Per 100K barcodes | | 100K vs 85K | 30 |
| Generate digital expression and plots for matched barcodes | 1 | 2 | 10 | | 4G | 50K | 110 |

### Supplementary Figures:

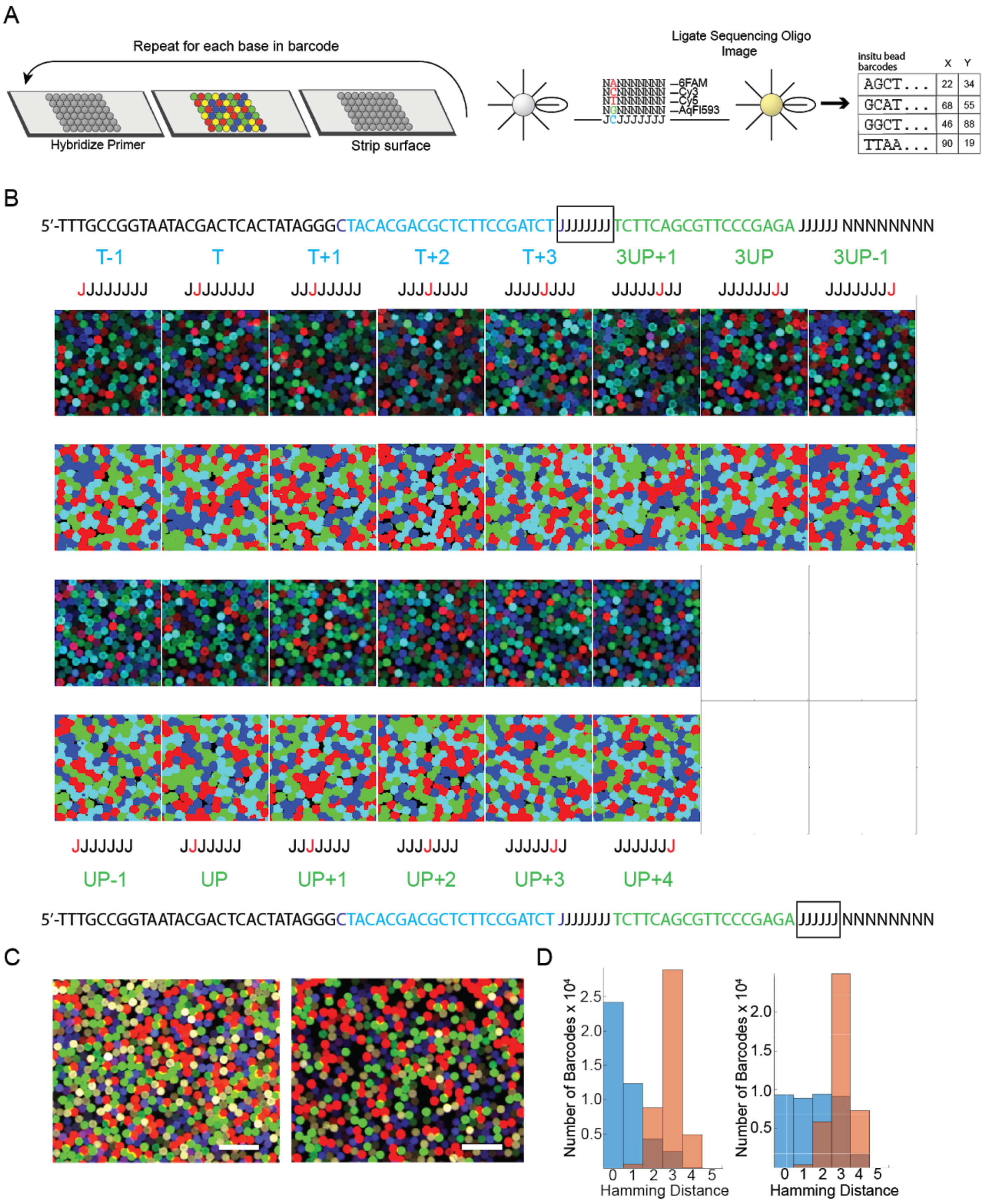

Figure S1: **Monobase sequencing by ligation chemistry allows for replacement of SOLiD sequencing for array generation**.

A) Overview of monobase sequencing scheme for indexing arrays.
B) Schematic representation of each base of monobase sequencing, and an example image as well as base-call for each base. Top: bases sequenced in the first block of Js using the Truseq (blue primers) as well as the 3’ sequencing off of UP (green primers). Bottom: bases sequenced in the second block of Js. Primer sequences are listed in Supp. Table 5. Red J base denotes J base being sequenced.
C) Lookup Table (LUT) matched images for monobase (left) and dibase (right) indexing.
D) Histogram of barcode matching (hamming distance to closest barcode) between imaging and short read sequencing data for array imaged in monobase (left) and dibase (right).

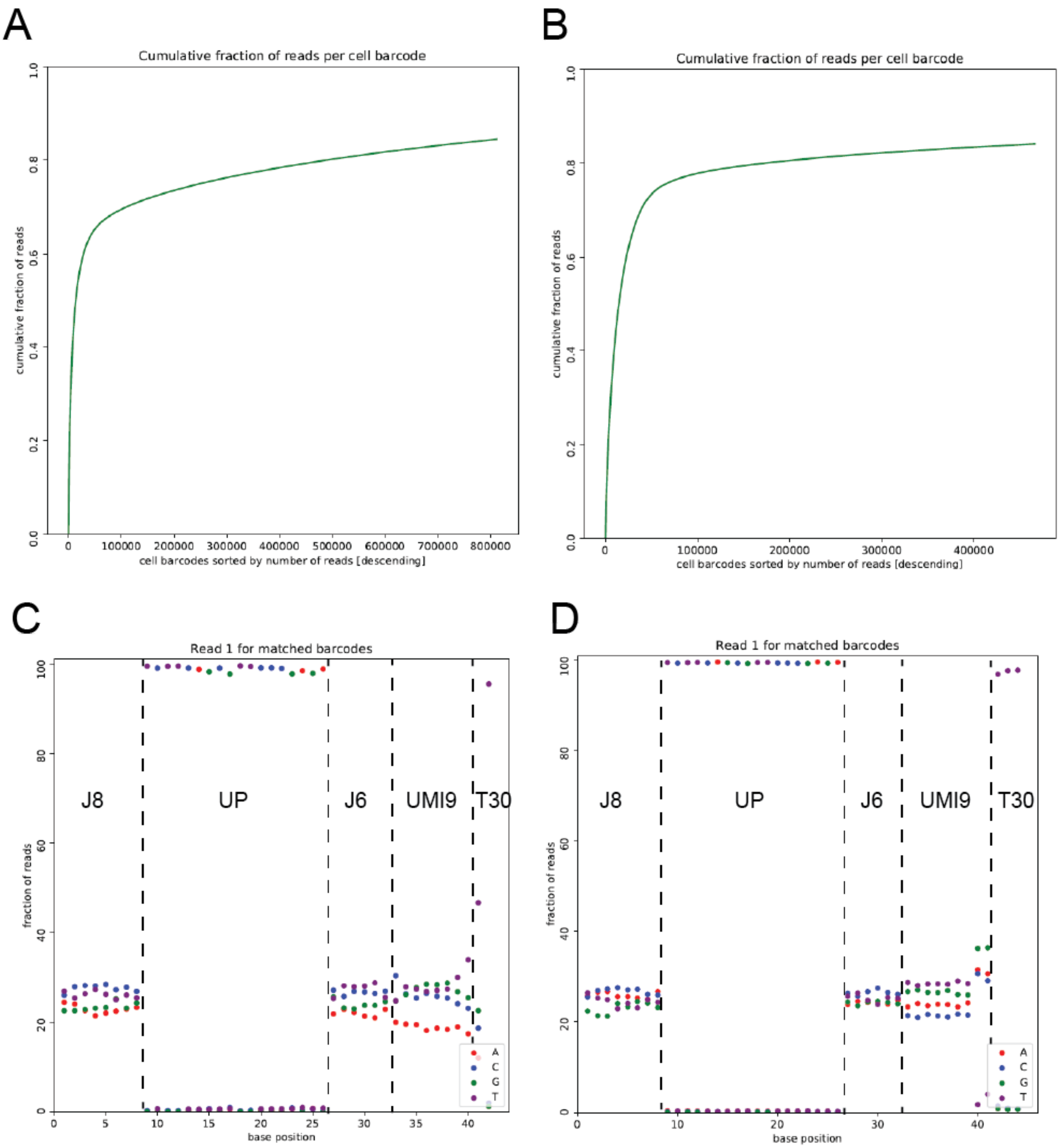


Figure S2: **Mapping and barcode matching statistics for barcoded beads used in Slide-seqV2.**

A) Cumulative distribution of reads mapping to barcodes for commercially supplied beads. There are ~100,000 beads sequenced, fraction of reads after 100,000 beads suggest synthesis errors resulting in fractured bead barcodes.
B) Cumulative distribution of reads mapping to barcodes for beads with custom optimized bead synthesis. There are ~100,000 beads sequenced, note the knee at 100,000 beads is ~80% of reads.

C) For barcodes that map to *in situ* sequenced beads, the base balance of the bead barcode for commercially supplied beads. Note the incomplete synthesis of bases in the constant bases of UP, as well as the high polyT% in the last base of the UMI, suggesting internal deletions in the bead sequence.

D) Same as C) but for custom optimized bead synthesis (vs2).

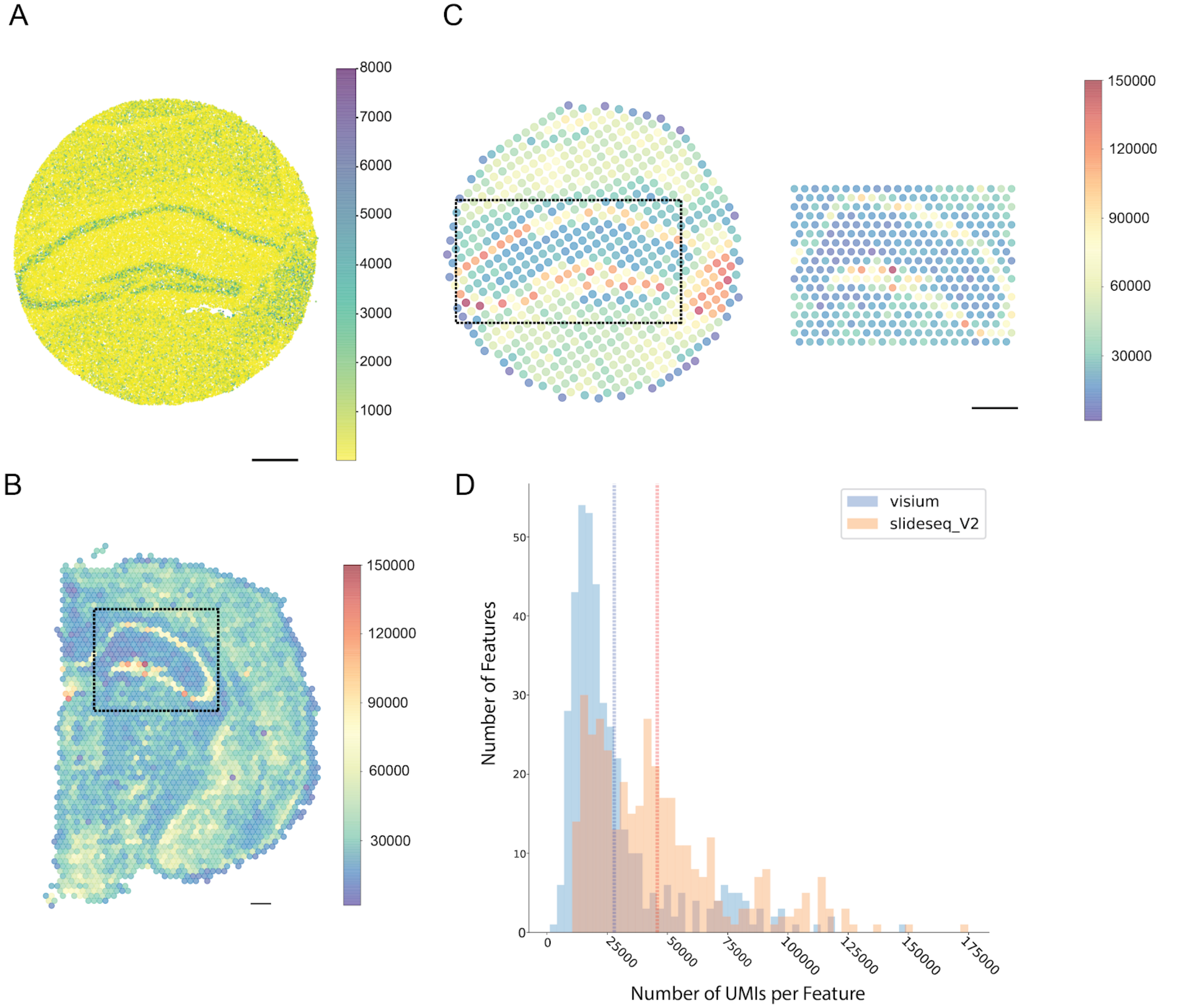


Figure S3: **Comparison of Slide-seq data to 10X Visium data.**

A) UMI counts of a Slide-seq experiment performed on the mouse hippocampus. Data displayed at 10μm resolution (Scale bar 500μm)

B) Visium data from the mouse hippocampus, scale bar represents number of UMIs.

C) Slide-seq data binned to equal size of the Visium data (110μm center to center for each spot). (Scale bars = 500μm). Scale bars represent the number of UMIs. Visium data chosen to match region from Slide-seq data (black box in B). Slide-seq data chosen for comparison surrounded by black box.
D) Histogram comparing number of spots per feature for binned Slide-seq data in C and Visium data. Dotted line represents the mean of each method. (Mean binned Slide-seqV2=45,772 UMIs, Mean Visium = 27,952 UMIs)

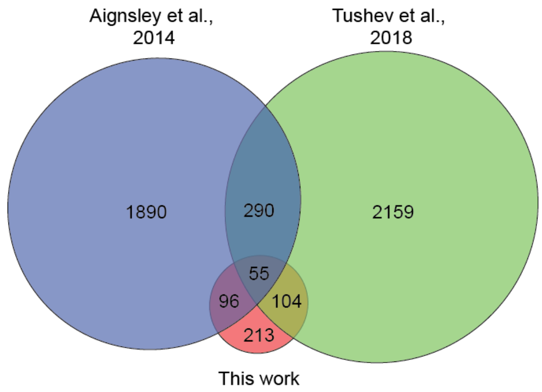


Figure S4: **Dendritically enriched RNAs identified in this work and two previous studies.** (Aingsley et al., 2014, and Tushev et al., 2018).

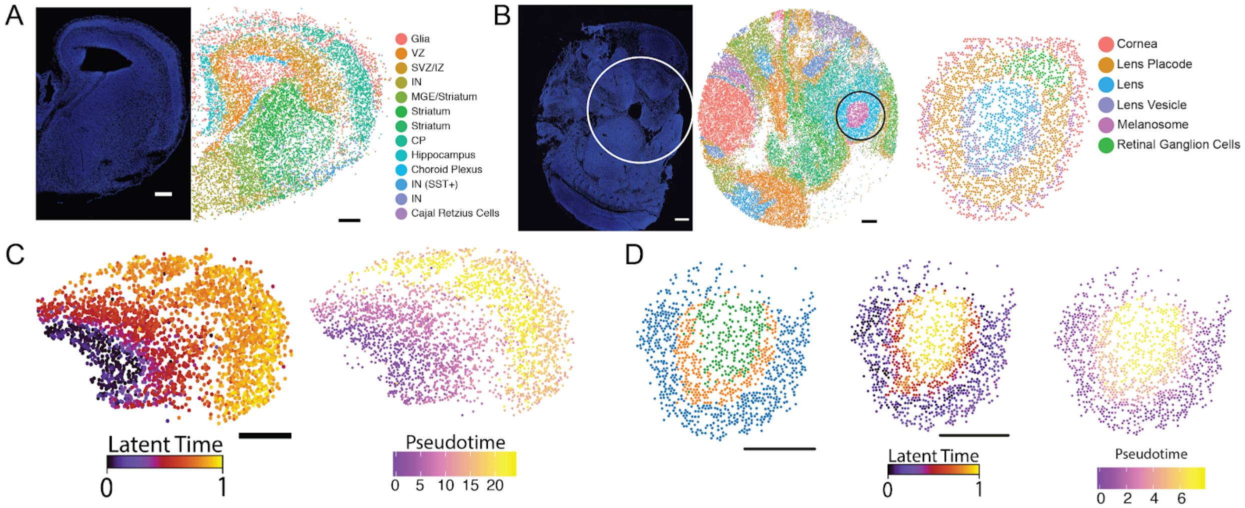


Figure S5: **Trajectory analyses of mouse E15 cortex and mouse E12.5 ocular lens.**
A) Left: DAPI stain of E15 mouse brain. This is a serial section of the region that was placed on the puck (scale bar 200µm). Right: Slide-seq data of E15 brain with cluster labels. VZ = Ventricular Zone, SVZ = Sub-Ventricular Zone, IZ = Intermediate Zone, IN = Interneuron, MGE = Medial Ganglionic Eminence, CP = Cortical Plate.
B) Section of E12.5 mouse embryo. Left: DAPI stained section with puck placement outlined in white line. Middle: Slide-seq data of embryo section. Coloring of beads by cluster. Black circle

represents the eye taken for downstream analysis. Right: Second round clustering of the eye with cluster labels representing different regions of developing eye.
C) Left: Latent time trajectory generated by scVelo on expression of Slide-seq for cortex data plotted in space (scale bar 200µm). Right: Pseudotime generated with Monocle3 plotted in space for cotex data.
D) Left: Developing lens taken for pseudotime analysis (Blue = Lens Placode, Orange= Lens Vesicle, Green = Lens). Middle: Same as scVelo figure in C but on lens. Right: Same as Monocle3 figure in C but on lens. (scale bars 200µm).

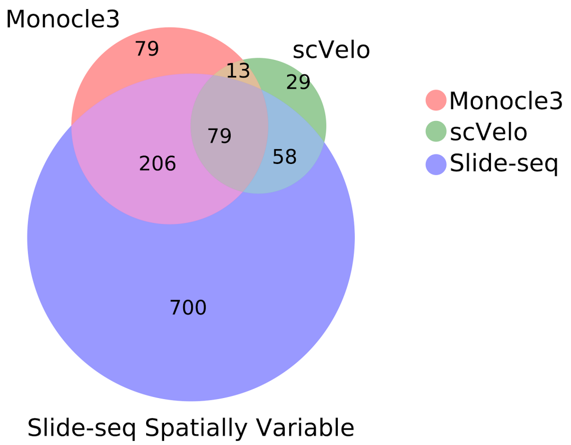


Figure S6: **Significant gene detection by trajectory inference methods versus Slide-seq**. Number of genes significantly loading onto Monocle3, scVelo, and the spatial developmental axis by Slide-seq in E15 mouse cortex.

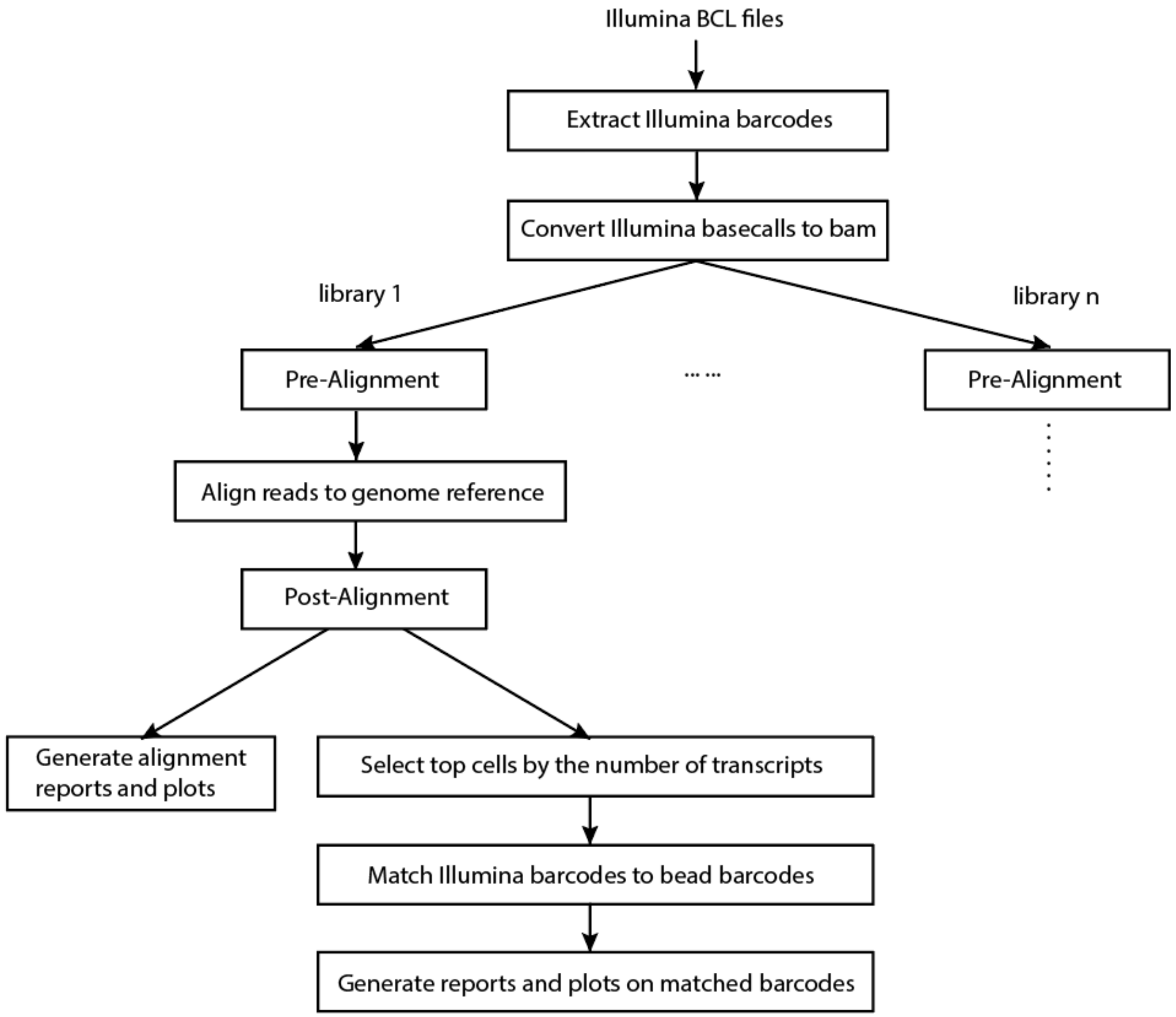


Figure S7: Workflow of the Slide-seq pipeline.
